# Impact of storage on the stability and the protective effect of extracellular vesicles released by *Candida albicans*

**DOI:** 10.1101/2025.09.13.675998

**Authors:** Leandro Honorato, Jhon J. Artunduaga Bonilla, Alessandro F. Valdez, Flavia C.G. dos Reis, Julio Kornetz, Albaniza Liuane Ribeiro do Nascimento Sabino, Marcio L. Rodrigues, Joshua D. Nosanchuk, Leonardo Nimrichter

## Abstract

Extracellular vesicles (EVs) released by *Candida albicans* are multi-antigenic compartments considered as promising prototypes for vaccine development. However, their stability, appropriate storage and handling conditions are largely unexplored, which raises questions related to their biotechnological applicability. Here, we evaluated the physical and functional stability of *C. albicans* EVs under long-term storage. Furthermore, we conducted a comparative analysis of these properties in *C. albicans* EVs obtained through three commonly utilized isolation protocols documented in the literature. After identifying the most efficient isolation method for optimal yield, we devised a potential quality control for EVs isolation based on protein and sterol ratio. Subsequently, we investigated the impact of drying EVs using vacuum centrifugation at room temperature or -4 °C and the effect of freeze-thaw cycles in EVs stability. Transmission electron microscopy (TEM) revealed that EVs maintained morphological stability after long-term (up to 4 years) storage at -80 °C as well as storage at room temperature, 4 °C and -20 °C for 7 days with or without vacuum centrifugation, with a tendency of higher recovery when lower temperature is used. Remarkably, all of the *C. albicans* EVs suspensions maintained their biological properties as demonstrated by their ability to protect *Galleria mellonella* against *C. albicans* infection. However, the number of freeze-thaw cycles significantly impacted on the protective effect of the EVs. Overall, our findings demonstrate that *C. albicans* EVs maintain notable morphological and biological stability of under several conditions, enabling their efficient and reproducible utilization in research and potentially as therapeutic agents.

**Importance:** Extracellular vesicles (EVs) released by *Candida albicans* are promising vaccine prototypes due to their multi-antigenic nature. However, their storage and handling conditions are not well understood, raising concerns about their biotechnological use. This study evaluated the long-term physical and functional stability of *C. albicans* EVs. We compared three isolation methods to identify the most effective one and suggested a quality control measure based on protein and sterol ratios. We also examined the effects of vacuum drying and freeze-thaw cycles on EV stability. Our findings show that *C. albicans* EVs maintain their biological function after long-term storage at -80 °C and under various conditions. Notably, their protective effect in an insect model was reduced though repeated freeze-thaw cycles. This research provides valuable insights for the efficient use of these vesicles in future studies.

## Introduction

Extracellular vesicles (EVs) are nanosized lipid bilayered compartments that carry a variety of biomolecules, characterizing a non-conventional secretion mechanism in virtually all cell types (1, 2). Despite being described only recently in comparison to other cell types, the EVs released by fungal organisms have been studied over the past 15 years. EVs from diverse fungal species contain a complex assortment of biomolecules associated with virulence, thus participating in multiple processes that can significantly impact the host-pathogen interactions (3–10).

*Candida albicans* EVs, specifically, have been extensively studied by our group and shown a promising potential as a vaccine platform. These EVs were demonstrated to effectively immunize insects and mice, protecting them against a subsequent lethal candidal infection (4–6, 11). More recently, we demonstrated that a combination of lipids carried by *C. albicans* EVs, including middle chain fatty acids and terpenes, are responsible for modulating yeast-to-hyphae differentiation, biofilm formation, and virulence, corroborating previous studies suggesting that fungal EVs are messenger compartments (12–14). EVs released by *C. albicans* biofilms are associated with both matrix formation and drug resistance (15, 16). However, the stability of vesicular components may poses challenges to harnessing the full potential of EVs. For instance, purified farnesol and dihydrofarnesol, proeminent terpenes carried by *C. albicans* EVs, gradually lose their potency to control dimorphism after storage under solution (14). This diminishing effectiveness prompts speculation that the microenvironment within EVs might play a role in preserving the stability of these components. In turn, it could lead to an extension of their half-life and consequently, enhancing the overall activities of the EVs. Thus, the functional integrity of EV compounds is pivotal, as any alterations or losses could undermine their biological activities and therapeutic efficacy. In a previous study, we demonstrated that EVs derived from *C. albicans* maintained their physical properties even after being stored for a week at temperatures of 4, -20 or -80 °C (4). Notably, these EVs retained their capacity to activate dendritic cells and provide protection against lethal candidiasis in *Galleria mellonella*, a wax moth model frequently used to investigate virulence, screening new drugs and study innate immune response (17, 18). However, EVs stability over longer storage periods have not yet been conducted.

In this study, our main goal was to examine the enduring physical and functional stability of *C. albicans* EVs preparations. We investigated this stability across four scenarios: (i) during long-term storage, (ii) utilizing three commonly used methods for fungal EV isolation, (iii) under vacuum centrifugation following variable temperature conditions and (iv) after freeze-thaw cycles. Our findings indicated that *C. albicans* EVs exhibit notable stability, rendering these compartments valuable as prototypes for biotechnological development.

## Materials and Methods

### Fungal strains and culture conditions

*C. albicans* strains ATCC 90028 and NGY 152 was stored in Sabouraud broth, 2% glucose (Sigma, USA) and 1% peptone (Acumedia, Brazil) with 15% glycerol (Merck, USA) and maintained at - 80 °C. Yeast cells were cultured in Sabouraud broth for 48 h at 30 °C under agitation (150 rpm).

### Preparation of fungal EVs

#### Ultrafiltration system (UF) system

EVs isolation was performed following a protocol established in our laboratory with few modifications (14). Briefly, cultures (one liter) were centrifuged at 4,000 x *g* for 15 min at 4 °C. The supernatants were collected and further centrifuged at 15,000 x *g* for 15 min at 4 °C in Beckman Avanti™ J-E to remove cell debris. Residual cells and debris were removed after a step of filtration using a 0.45 μm membrane filter (Merck Millipore, USA). The cell-free supernatant was concentrated about 50 times using an Amicon® stirred cells system (100 kDa membrane, Merck Millipore, USA). The concentrated supernatant (20 ml) was then centrifuged at 100,000 x *g* for 1 h at 4 °C in Beckman Optima™ LE-80K, rotor 70ti (k-factor = 156). The pellet resultant was washed twice with phosphate-buffered saline (PBS) pH 7.4, at 100,000 × *g* for 1 h at 4 °C.

#### Tangencial flow ultrafiltration (TFUF) system

EVs isolation was adapted from a protocol established by Heniemann and colleagues (19). The cell-free supernatant was obtained under the same conditions described above for the ultrafiltration system and then concentrated 50 times using an Tangential Flow Filtration System (VivaFlow 200 flipflow filtration MWCO 100 kDa Sartorius, USA). To concentrate the supernatant, the Masterflex easy-load II (model 77200-60, USA) pump was used at speed 4, flow rate of 25 ml/min. The concentrated supernatant (approximately 20 ml) was then centrifuged at 100,000 x *g* for 1 h at 4 °C in Beckman Optima™ LE-80K. The pellet resultant was washed twice with PBS pH 7.4, at 100,000 × *g* for 1 h at 4 °C. The number of cells at the end of the culture was determined by counting them using a Neubauer chamber.

#### EV isolation from solid media (SM)

EVs isolation was performed following a protocol established by Reis *et al* (20), Briefly, a cell suspension was made with 3 x10^7^ cells/ml and 300 μL, containing approximately 10^7^ yeasts of *C. albicans* yeast cells, were plate onto Petri dishes containing Sabouraud Dextrose Agar (SDA). After 24 hs of growth, cells were harvested using a cell scraper and transferred to 20 ml of PBS. The cell suspension was centrifuged at 4,000 x *g* for 15 min at 4 °C. The supernatants were collected and further centrifuged at 15,000 x *g* for 15 min at 4 °C in Beckman Avanti™ J-E to remove cell debris. Residual cells and debris were removed after a step of filtration using a 0.45 μm membrane filter (Merck Millipore, USA). The concentrated supernatant (20 ml) was then centrifuged at 100,000 x *g* for 1 h at 4 °C in Beckman Optima™ LE-80K. The pellet resultant was washed twice with PBS pH 7.4, at 100,000 × *g* for 1 h at 4 °C. All fungal EVs were then suspended in PBS and aliquots were plated onto brain heart infusion (BHI) agar (Sigma, USA) plates and incubated for 72 h to confirm the absence of *Candida* contamination.

#### EVs protein quantification

The quantification of EVs was carried out using the bicinchonininc acid (BCA) Protein Assay Kit (ThermoFisher, USA) following the manufacturer’s instructions.

#### EVs sterol quantification

The assessment of EV fractions was quantified by measuring the presence of sterols, employing the Amplex Red Sterol Assay Kit (Molecular Probes, Life Technologies) through a quantitative fluorimetric method (6).

#### Electron Microscopy

EVs morphology and size were assessed using transmission electron microscopy (TEM) employing the negative staining technique. In summary, EVs (5 μl) were applied onto a Formvar-coated, carbon-coated 300-mesh copper grid and allowed to adsorb for 30 seconds. Excess solution was removed using filter paper, and then the samples were stained with 2.5% uranyl acetate for an additional 30 seconds. EVs micrographs were captured using a transmission electron microscope (FEI Tecnai Spirit) operating at 120 kV. The diameter of EVs was measured using ImageJ software (version 1.53k). Up to 150 EVs per sample were analyzed, based on observations from up to 10 micrographs per sample. EVs from up to two replicates were examined. The data obtained from ImageJ were used to generate relative frequency distribution histograms in GraphPad Prism 8.0 software.

#### Freeze-thaw cycles

For the freeze-thaw (FT) cycles, EVs isolated from the tangential flow ultrafiltration system were separated into 20 μl aliquots with a concentration of 1 mg/ml based on protein content. Aliquots were thawed and frozen 2, 4, 6, 8 or 10 times at -80 °C in the presence or absence of cryopreservative (10% sorbitol). To remove the sorbitol the EVs suspensions were ultracentrifuged at 100.000 x *g* for 1 h and the supernatant was discarded.

Concentration of protein and sterol were verified as described above.

#### Galleria mellonella infection

*G. mellonella* larvae in the final instar larval stage were selected according to similarity in weight (0.3—0.35 g). Larvae (10 per group) were inoculated with 10 μl of EV suspensions (100 μg/ml per insect, based on protein quantification) using a Hamilton syringe into the haemocoel through the last proleg. The same volume of PBS was used as negative control in different larva from the same collection of *Galleria*. The larvae were then placed in sterile Petri dishes and kept in the dark at 37 °C for two days. Subsequently, all larvae were inoculated with 10 μl of a suspension containing 2 × 10^5^ yeasts of *C. albicans* (ATCC 90028). Larvae were kept under the same conditions as above and they were monitored twice daily for survival. Death was determined by the lack of movement in response to physical stimulation. Survival curves were plotted, and statistical analyses were performed using the log-rank (Mantel–Cox) survival test.

#### Statistical analysis

All statistical analyses were performed using: one-way ANOVA and analyzed by Dunnett’s multiple comparisons test; for survival, and the difference between groups was analyzed by log-rank (Mantel–Cox) test were performed with the GraphPad Prism 6, version 6.02 for Windows (GraphPad Software). The dataset containing the statistical analyses is available at the following link https://data.mendeley.com/preview/kjhffkf7c2?a=fae7e0c5-d6aa-42b1-8360-042e4c4b0871.

## Results

### Long-term storage modified *Candida albicans* EVs size

For long-term storage experiments, all EVs were collected using the ultrafiltration system (6) and then stored in PBS at -80 °C. Samples were processed and TEM was used to determine EVs size and integrity and the data compared with a fresh preparation of EVs. The majority of EVs from the different conditions displayed round and cup-shaped compartments, ranging from 30 to 100 nm. However, EVs stored for over three years had a small but significative increase in their measurements. The frequency of EVs stored for 3 and 4 years within the 30-100 nm range decreased and more EVs with higher diameters were observed (Fig. 1). EVs after 2 years of storage displayed an intermediate profile, with a higher percentage of smaller EVs (between 30 and 90 nm), but distinct from 1 year storage and fresh EVs.

**Figure 1.**
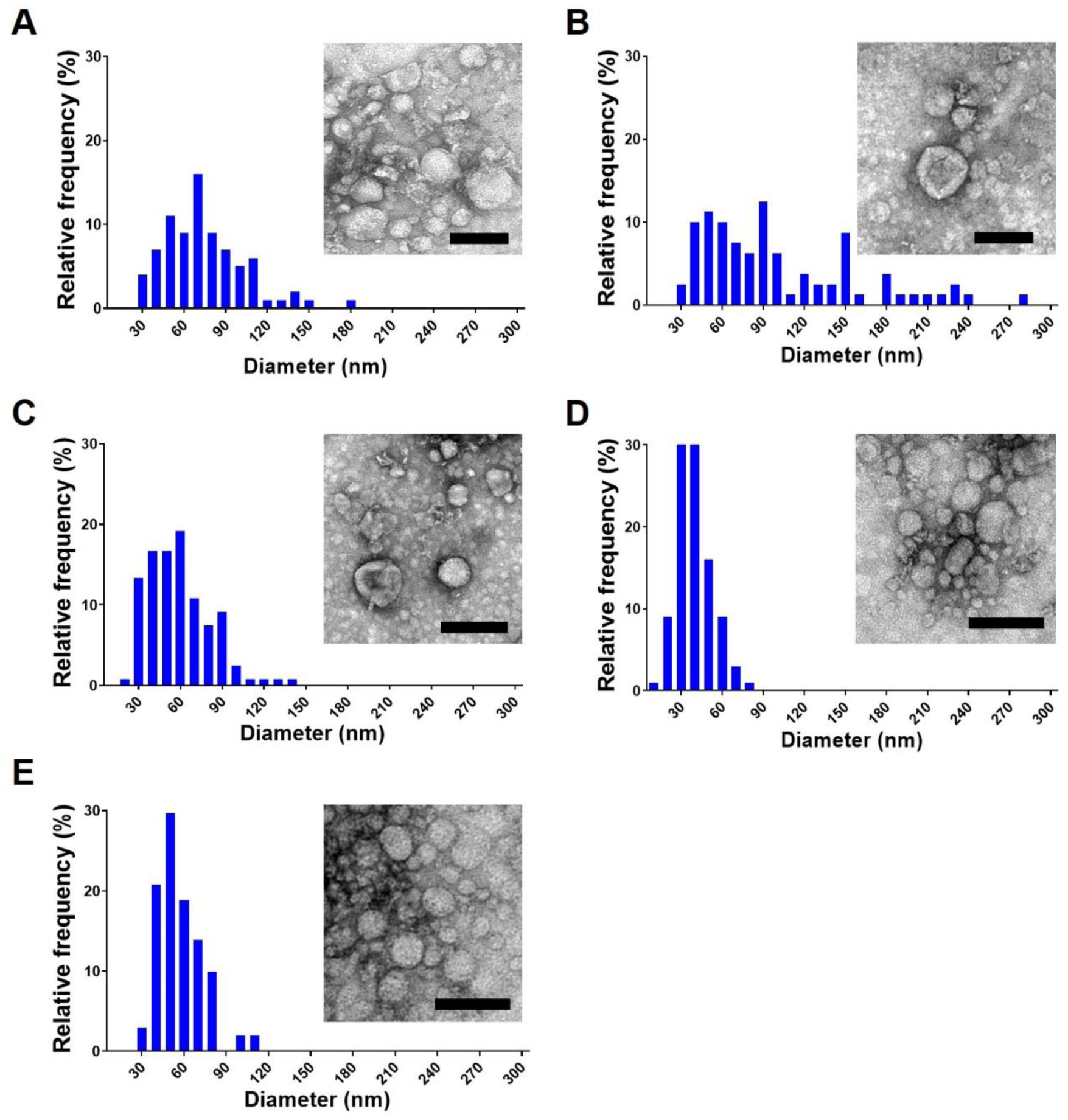
Effect of storage time on the dimensions and morphological properties of *Candida albicans* EVs. The stability of *C. albicans* ATCC 90028 EVs was studied over a period of 4 years. Vesicles stored for four years (A), three years (B), two years (C), and one year (D) were visualized and compared with fresh EVs (E) using TEM after negative contrast staining with uranyl acetate. The histograms in A, B, C, D and E depict the size distribution of EVs after measuring 100 EV/sample using ImageJ software. The mean diameter of the EVs were compared using Tukey’s multiple comparisons. Scale bars, 100 nm.

### EVs retain their protective effects in *G. mellonella* independently from the duration of storage

We found that EVs submitted to variable storage times maintained their ability to protect *G. mellonella* larvae against lethal *C. albicans* infections. Although animals pre-inoculated with PBS (control) succumbed to infection within seven days, EV-inoculated animals exhibited protection levels ranging from 70% to 90% (Figure 2). Although fresh EVs exhibited the highest protection percentage, the differences between the different years of storage were not statistically significant.

**Figure 2.**
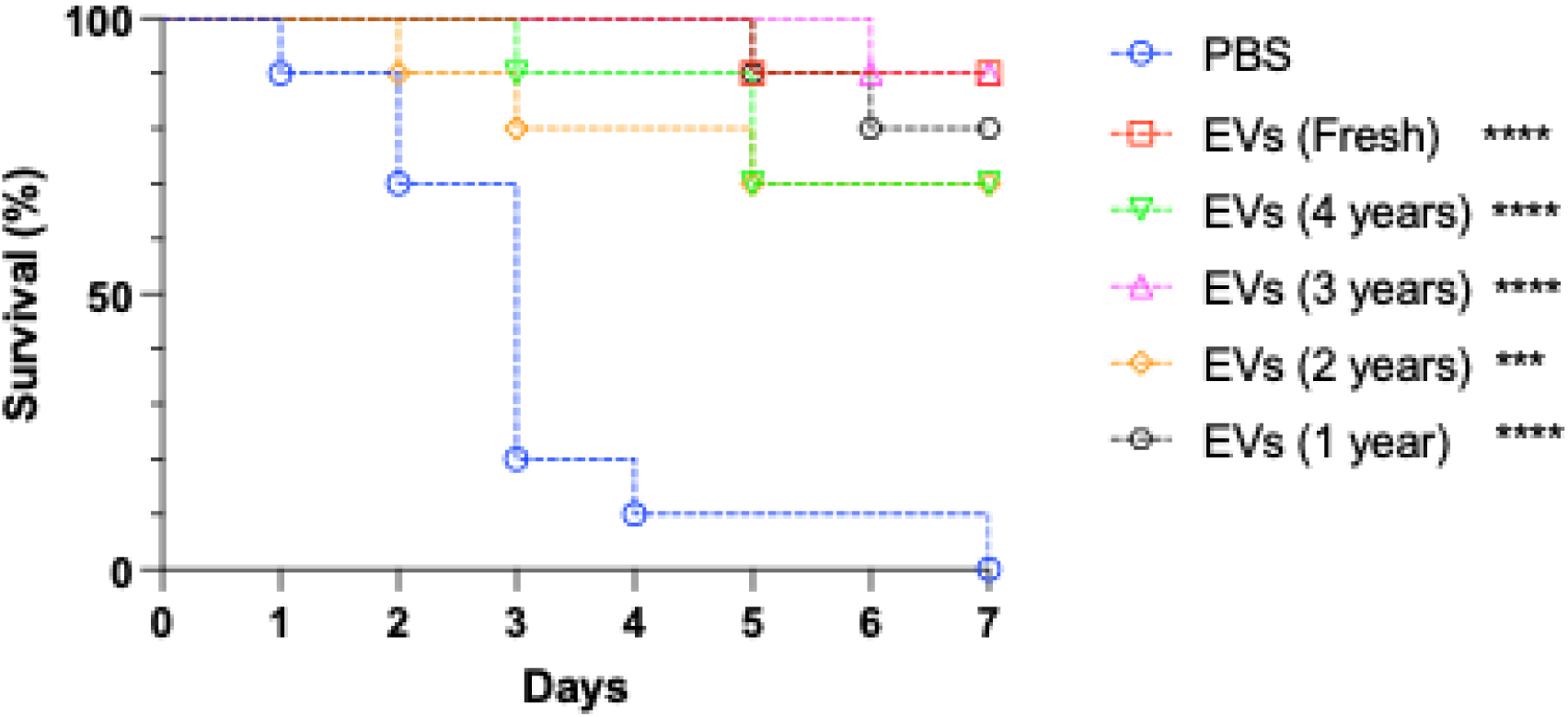
– Protection of *G. mellonella* larvae against *C. albicans* by EVs stored for different periods of time. Larvae from *G. mellonella* were inoculated with *C. albicans* (strain 90028) fresh EVs suspensions or EVs obtained using the Amicon system that were stored for different lengths of time. A final volume of 10 μl of a 100 μg/ml EVs suspension, based on the protein content, was injected. PBS was used as control (absence of EVs). The insects were infected two days later with a lethal inoculum of *C. albicans* (2 x 10^5^ cels) (ATCC 90028). The survival differences between each EV groups and PBS were analyzed by individual paired Log-rank (Mantel-Cox) **** p <0.0001, *** p= 0.0001. Mortality was monitored for 7 days (n=10). Results are representative of two independent experiments.

### Culture conditions and isolation procedure impact EVs size and time consuming

To evaluate whether the protocol used for EV isolation affects their properties and protective effect, we used three different approaches for vesicle preparation (Fig. 3). Despite generating EVs with comparable characteristics, there is notable variation in their sizes according to the protocol used. EVs obtained from liquid cultures (Fig. 3A and B, respectively) manifested dimensions ranging mostly from 20 to 100 nm (Fig. 3A and B). In contrast, EVs isolated from solid medium displayed a size distribution of 30 to 400 nm (Fig 4C).

**Figure 3.**
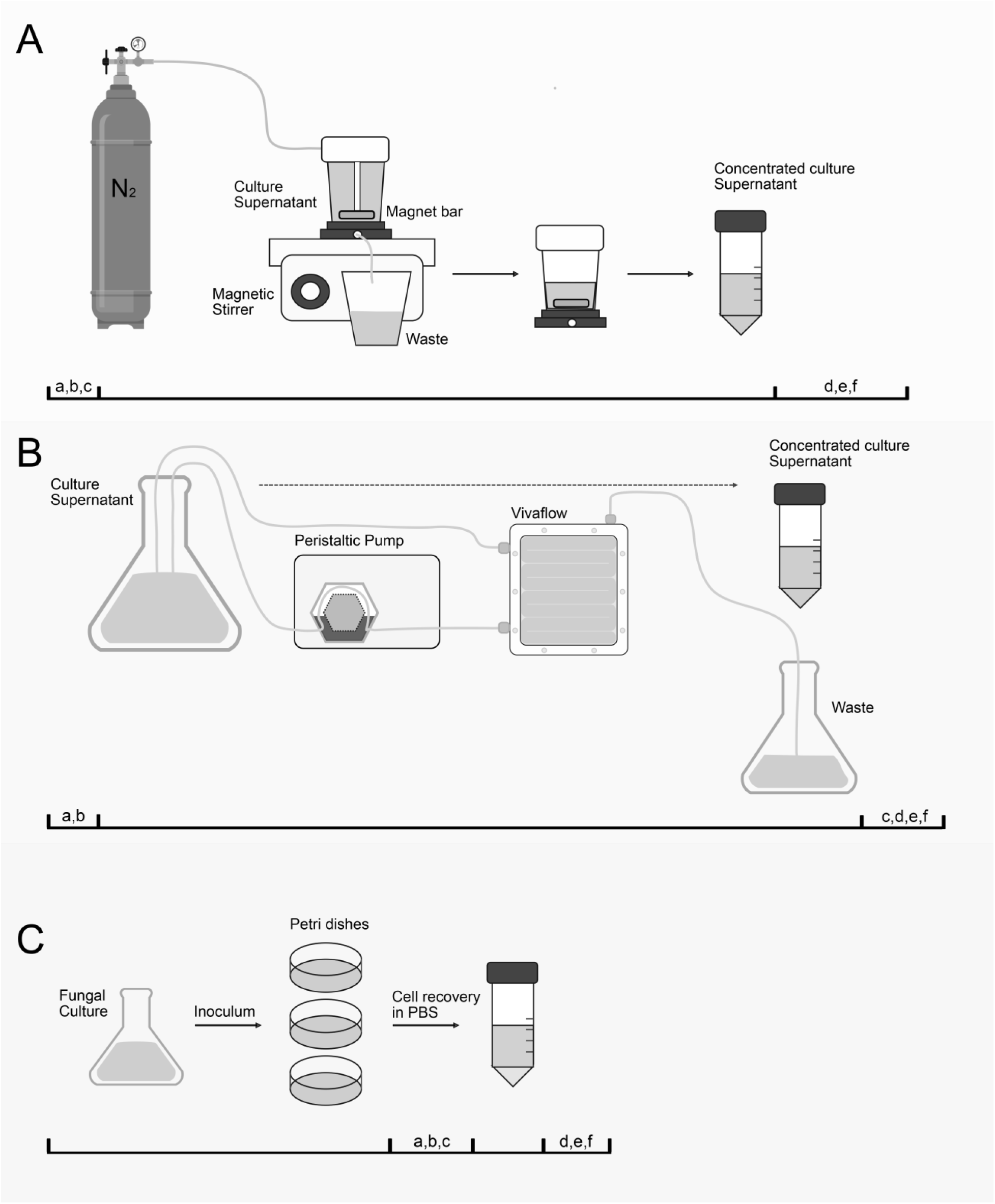
Schematic illustration depicting the three most common procedures used to isolate EVs released by fungal organisms. EVs isolation using an ultrafiltration system (A), a tangential ultrafiltration system (B) and solid medium (C). A list of common steps include: a) centrifugation at 4,000 x *g* for 15 min at 4 °C (Beckman Avanti^TM^ J-E), b) centrifugation at 15,000 x *g* for 15 min at 4 °C (Beckman Avanti^TM^ J-E), c) filtration using a 0.45 μm membrane filter (Merck Millipore, US), d) centrifugation at 100,000 x *g* for 60 min at 4 °C (Beckman Optima^TM^ LE-80K), e) quality control: EVs are suspended in PBS, plated on BHI agar, and left to incubate for 72 h (the absence of *Candida* colonies confirm purity of EV preparation, and f) EVs quantification using bicinchonininc acid (BCA) Protein Assay Kit and/or Amplex^TM^ Red cholesterol kit (Thermofischer, US). The estimated time for concentrating one liter of supernatant for ultrafiltration system and tangential ultrafiltration system are approximately 6 h or 30 min, respectively.

**Figure 4.**
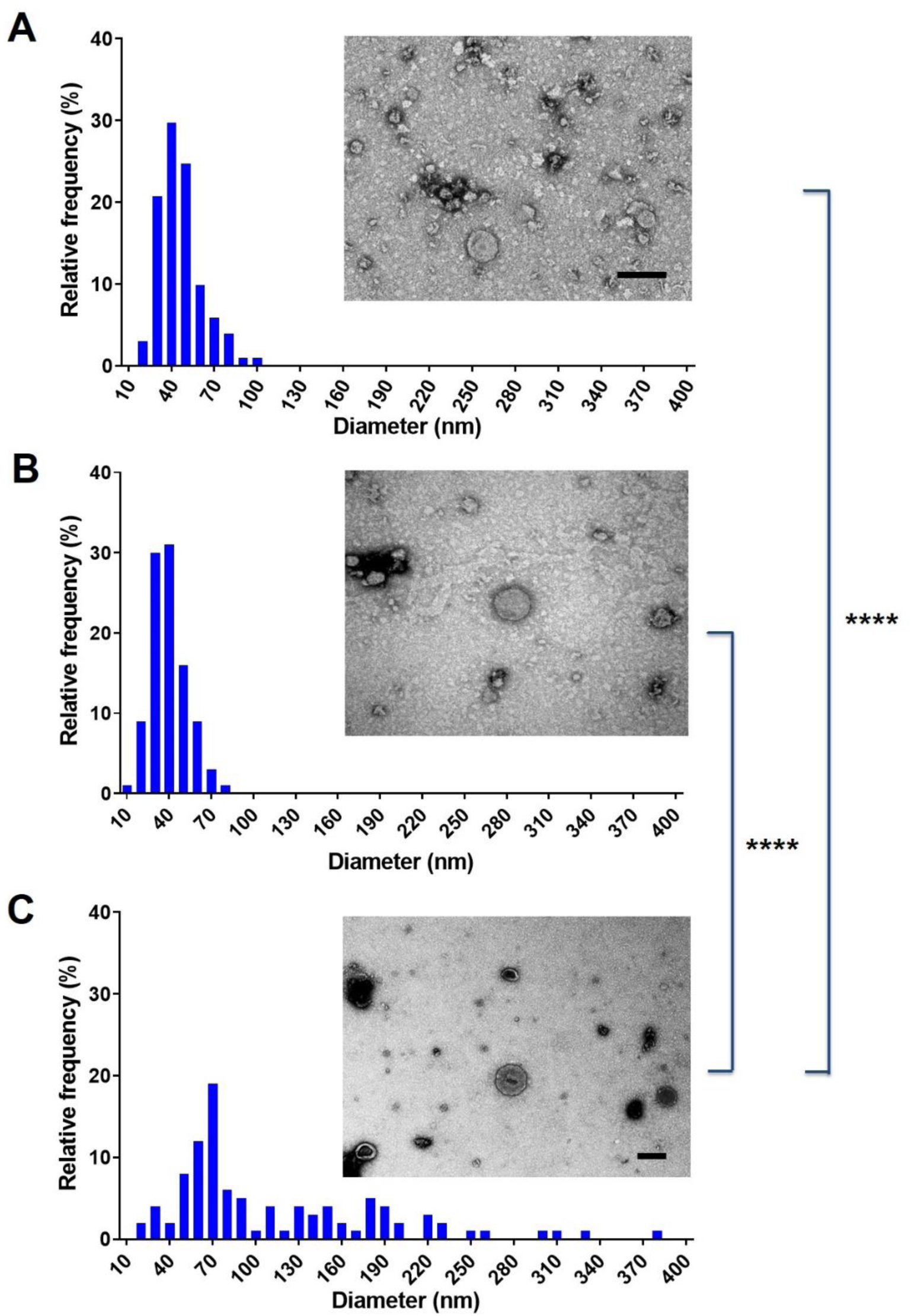
Comparison of the size distribution of EVs isolated from liquid or solid medium. *C. albicans* ATCC 90028 EVs were concentrated separately from liquid medium using a tangential ultrafiltration system (UF), flow ultrafiltration system (TFUF), or from solid medium (SM) on Petri dish. The diameter distribution of EVs obtained by TFUF (A), UF (B) and from SM (C) was determined using ImageJ software by measuring 100 EV/sample. EVs were visualized by TEM (inset images). Scale bar in insets represents 50 nm. **** p<0.0001.

### The protective effect of fungal EVs was not affected by the isolation method

To assess and compare the protective effects of fresh EVs obtained through the three distinct methods, we once again employed *G. mellonella* as a model organism. As depicted in Figure 5, all insects succumbed to the *Candida* infection within a week under control conditions (PBS injected larvae). However, regardless of the EV isolation method utilized, larvae pre-treated with EVs exhibited resistance to *C. albicans* infection. While EVs obtained from liquid medium were more effective, providing complete protection, there were no statistically significant differences between the protective effects of EVs isolated using ultrafiltration system (UF) or tangential flow ultrafiltration (TFUF) when compared to EVs derived from solid medium (SM).

**Figure 5.**
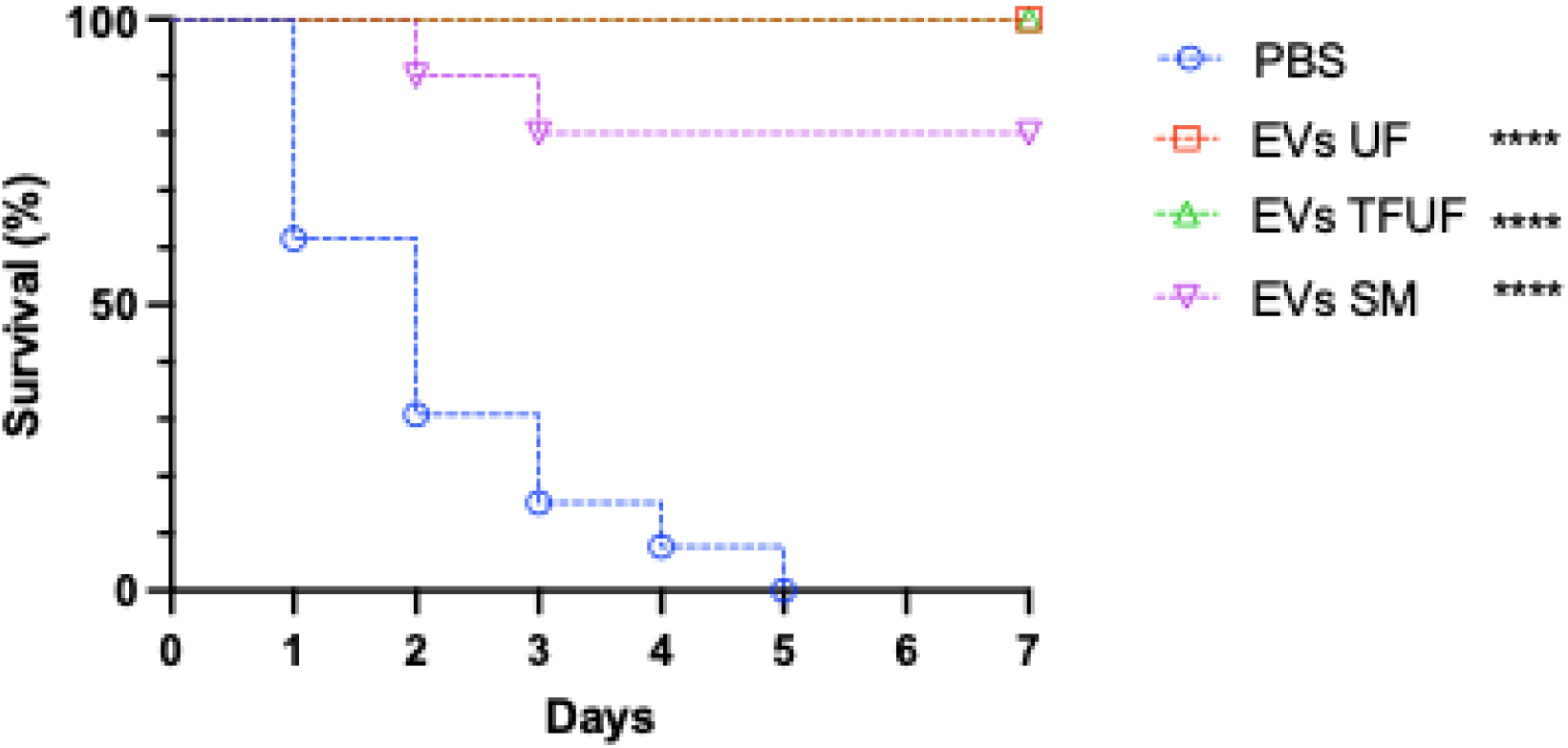
Protective effect of EVs isolated using distinct protocols for EV isolation. Survival curves of larvae pretreated with EVs isolated using Ultrafiltration system (UF, liquid medium), Tangential flow Ultrafiltration (TFUF, liquid medium) or solid medium culture (SM). Larvae from *G. mellonella* were inoculated with the distinct EVs suspensions (10 μl of a 100 μg/ml suspension based on the protein content). PBS was used as control. EVs were stored at -80 °C for up to 7 days before use. All *G. mellonella* larvae were infected two days later with a lethal inoculum of *C. albicans* (2 x 10^5^ cels) (ATCC 90028) yeasts are shown. Survival was monitored for 7 days (n=10). The survival differences between each EV group and PBS were analyzed by individual paired Log-rank (Mantel-Cox) **** *p* <0.0001. Results are representative of two independent experiments.

### Establishing sterol and protein ratio as a step for quality control of EVs isolated by *C. albicans* strains

Ensuring the quality of EV isolation poses a significant challenge, demanding the ability to discern between successful and suboptimal procedures. To address this, the overall sterol and protein contents were normalized based on the number of yeast cells, allowing for an initial comparison of EVs from two distinct *C. albicans* strains (Fig. 6A-C). Upon plotting various EV preparations, we found a consistently high protein-to-sterol correlation coefficient exceeding 0.98 for all isolated EVs (Fig. 6D). This robust correlation underscores the reliability of our methodology across diverse EV samples, offering a valuable indicator of the quality of isolation procedure.

**Figure 6.**
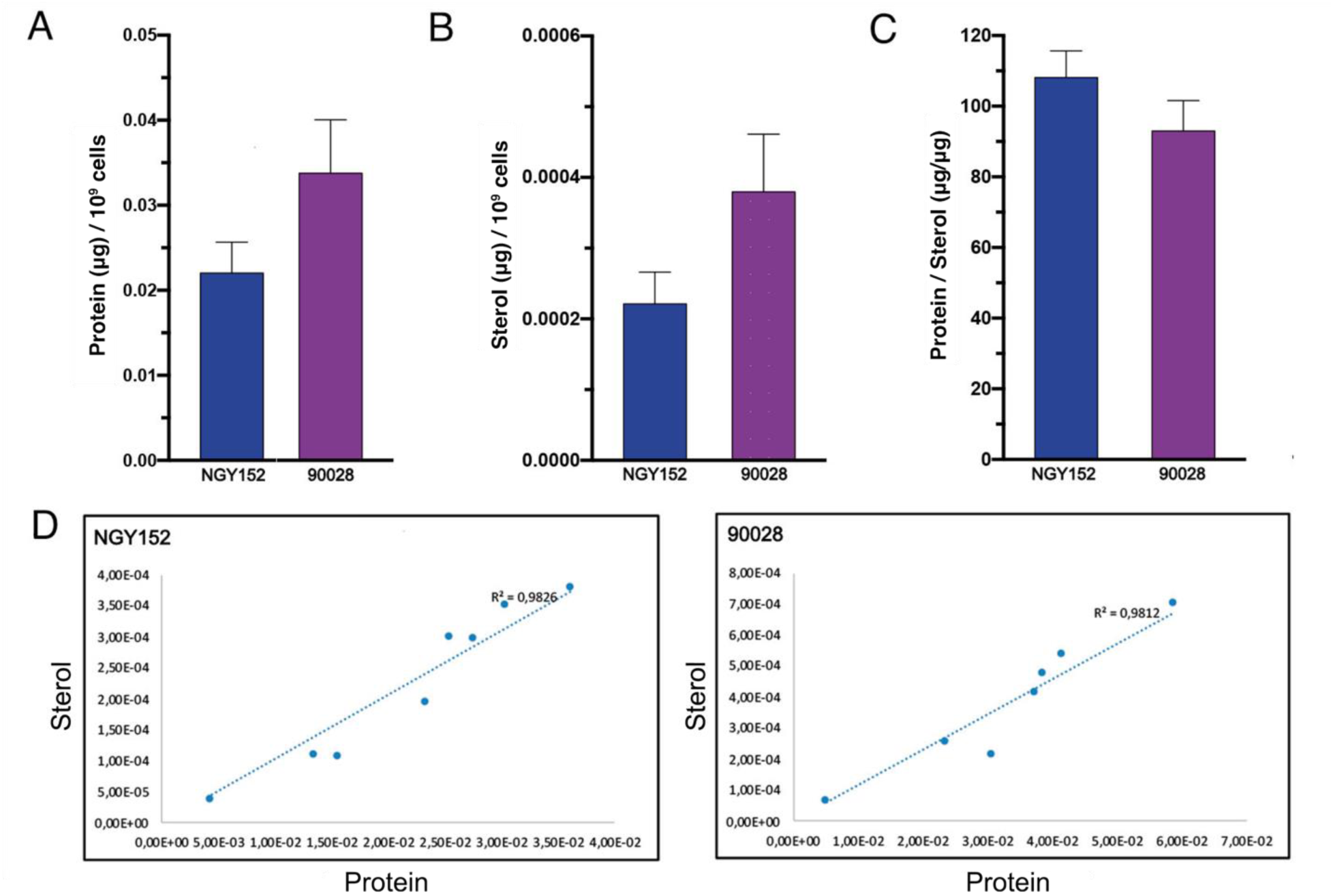
Protein and sterol ratio in EVs released by *C. albicans* strains. EVs were isolated from *C. albicans* strains NGY152 and 90028. The amounts of protein and sterol of *C. albicans* EVs were normalized by the number of yeast cells, enumerated using a Neubauer chamber at the end of the culture (A and B) and the protein to sterol ratio calculated (C). The correlation curve for EVs from each fungal strain is shown (D). Error bars represent standard deviation. Results represent the average of 7 independent EVs isolations. Statistical analysis were performed using one-way ANOVA and analyzed by Dunnett’s multiple comparisons test. No differences were found when the protein/sterol ratio was compared.

### Vacuum centrifugal evaporation of EVs does not compromise their biological activities

Due to its reduced time demands, the tangential flow ultrafiltration (TFUF) system was established as the optimal protocol for isolating EVs from large volumes of supernatant. Using EVs isolated from larger volumes (4 liters), we examined the physical stability of *C. albicans* EVs under various storage conditions, including vacuum evaporation concentration at room temperature (VCRT) and at -4 °C (VCLT), and compared them with control conditions (fresh EVs stored for a week in PBS at -80 °C). TEM data revealed that irrespective of the storage conditions, all samples exhibited EV morphologies similar to the control (Fig. 7) or as observed in freshly isolated EVs (see Fig. 4A). There were minor differences noted based on the drying condition, with VCRT displaying slightly larger diameters, suggesting the fusion of some particles (Figure 7).

**Figure 7.**
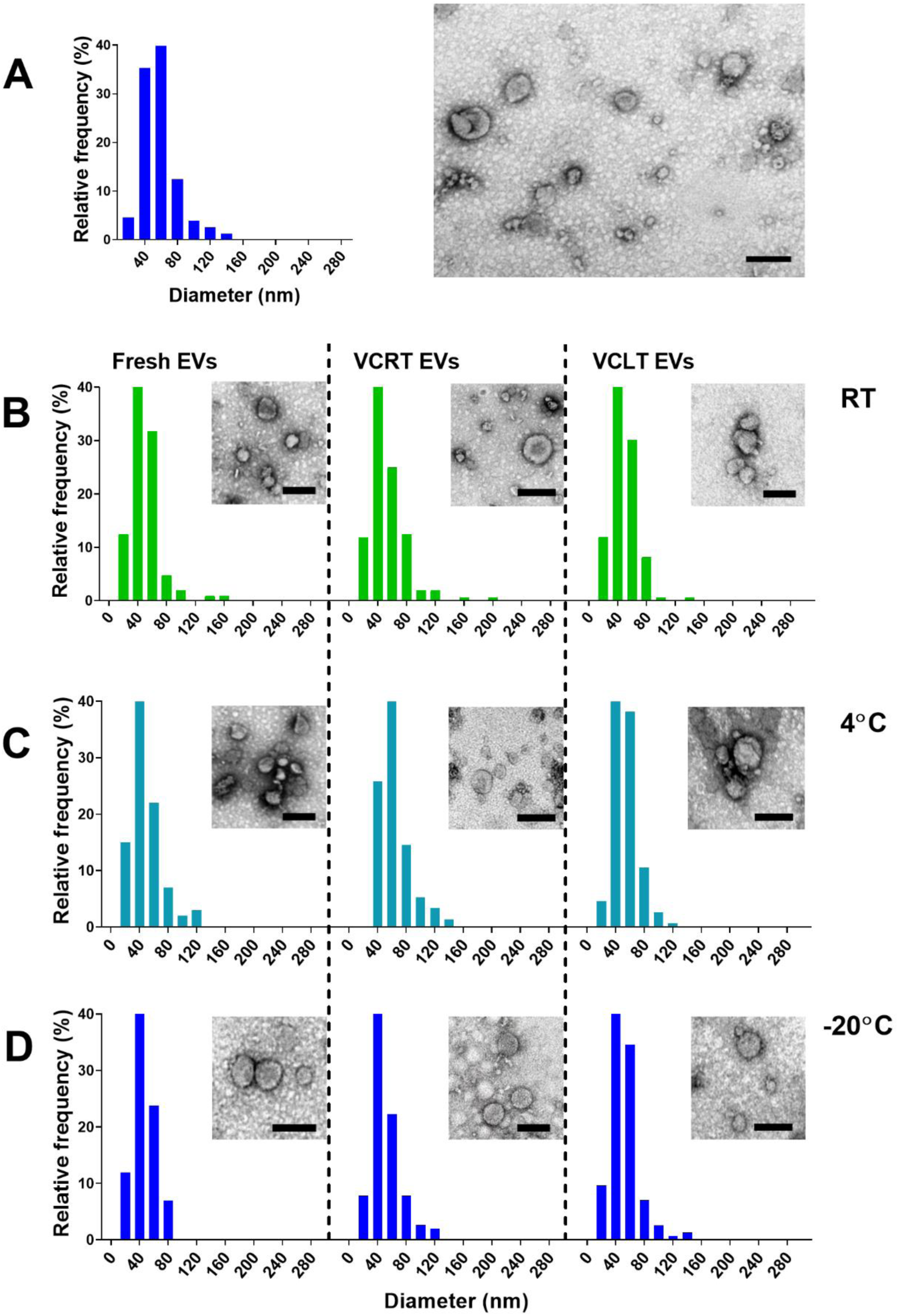
Effects of vacuum centrifugation on the stability and morphological properties of *Candida albicans* EVs. *C. albicans* ATCC 90028 vesicles were isolated using the tangential flow protocol. Fresh suspensions of EVs (A) and dried preparations obtained through vacuum centrifugation at room temperature (VCRT-EVs), or at -4 ⁰C (VCLT-EVs) were separately stored at room temperature (B), 4 ⁰C (C) or −20 ⁰C (D) for 7 days. EVs were suspended in PBS, negatively stained with contrast and then visualized by TEM. The frequency distribution of EVs diameter stored at RT, 4 ⁰C and −20 ⁰C, was determined by the ImageJ software at 150 EV/sample. Scale bars, 100 nm.

### Impact of ultracentrifugation and detergent treatment on EV recovery

To assess the impact of different treatments on *C. albicans* EVs, we first evaluated the loss caused by a single ultracentrifugation (UC) step. Our results showed that an additional UC run decreased the amounts of proteins and sterols by approximately 20% (Fig. 8). Only minor differences were observed when EVs were stored at XX °C for 4 h. In contrast, treatment with Triton X for 4 h reduced EV recovery to ∼50%. After 24 h in the presence of Triton X, no protein was detected and only trace amounts of sterols remained, indicating that the EVs were almost completely disrupted.

**Figure 8.**
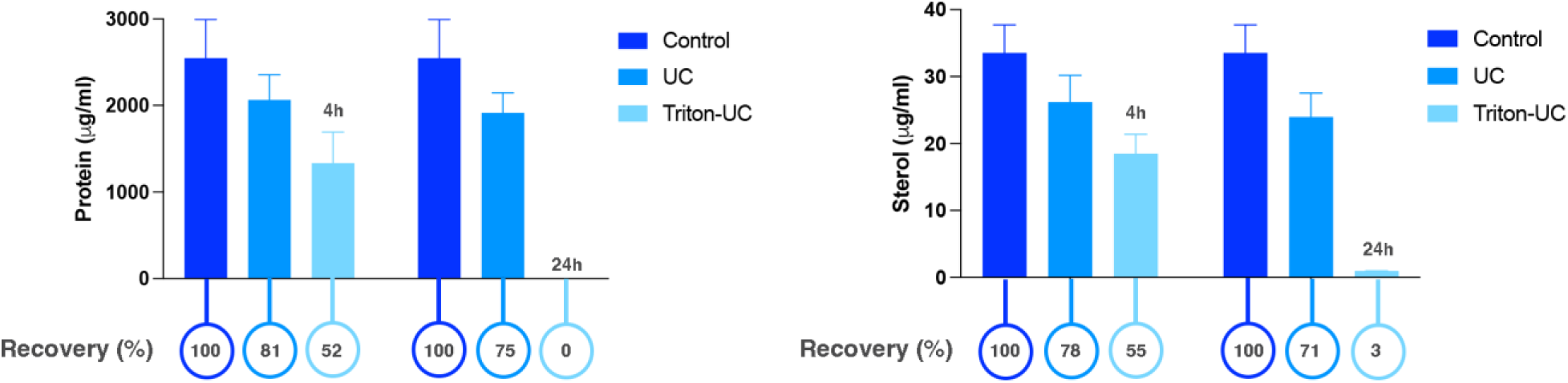
Recovery of EVs from *Candida albicans* after one UC step. *C. albicans* ATCC 90028 vesicles were isolated using the tangential flow protocol. Fresh suspensions of EVs were storage for 4 (A) and 24 h (B) with or without the addition of trition-X (X%). Subsequently, the EVs suspensions were submitted to an additional step of UC and their protein and sterol content determined (circles below the graphic). The percentual recovery of EVs was determined. No statistical differences were observed.

### Comparable reduction in protein and sterol content suggests loss of EVs across storage conditions

The recovery analysis of *C. albicans* EVs, determined indirectly as protein and sterol content, revealed similar trends across all storage and processing conditions, although no statistically significant differences were detected (Fig 9). In all cases, storage at –20 °C tended to yield higher recovery values, while storage at 4 °C showed lower recovery specifically when the EVs were kept *in natura.* Notably, protein and sterol measurements exhibited highly similar patterns of reduction, supporting the interpretation that the observed decreases reflect an overall loss of EVs rather than selective degradation of individual components.

**Figure 9.**
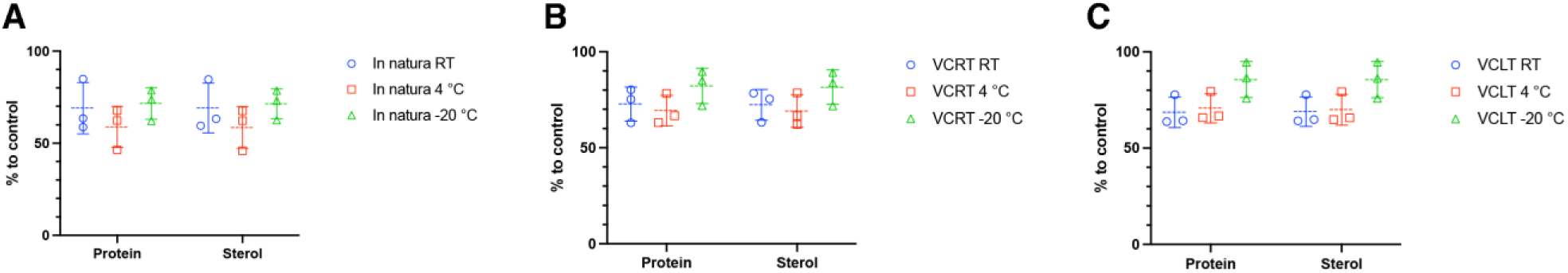
Effects of vacuum centrifugation on the recovery of *Candida albicans* EVs. *C. albicans* ATCC 90028 vesicles were isolated using the tangential flow protocol. Fresh suspensions of EVs (A) and dried preparations obtained through vacuum centrifugation at room temperature (VCRT-EVs), or at -4 ⁰C (VCLT-EVs) were separately stored at room temperature (B), 4 ⁰C (C) or −20 ⁰C (D) for 7 days. Protein and sterol quantification were performed and the recovery of the EVs were measured and compared between the distinct storage conditions. No statistical differences were observed.

**Figure 10.**
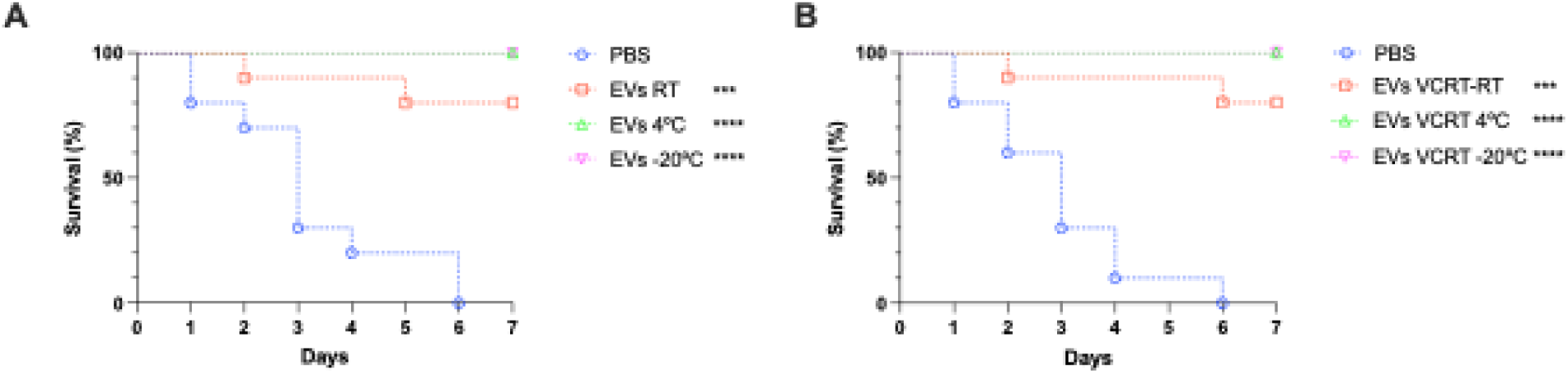
– Protective effect of vacuum concentrated *Candida albicans* EVs. *C. albicans* ATCC 90028 EVs were obtained using the Vivaflow System. Control or vacuum centrifugal evaporated EVs (room temperature, RT) were stored at -20 °C, 4 °C or RT for a week. Survival curves of *G. mellonella* larvae pretreated with *in natura* (A) or vacuum concentrated rehydrated (B) EVs (10 μl of a 10 μg/ml suspension based on the protein content). PBS was used as control. Larvae were infected two days later with a lethal inoculum of *C. albicans* yeasts (2 x 10^5^ cels) (ATCC 90028) are shown. Survival was monitored for 7 days (n=10). The survival differences between each EV groups and PBS were analyzed by individual paired Log-rank (Mantel-Cox) **** p <0.0001, *** p= 0.0001. Results are representative of two independent experiments.

**Figure 11.**
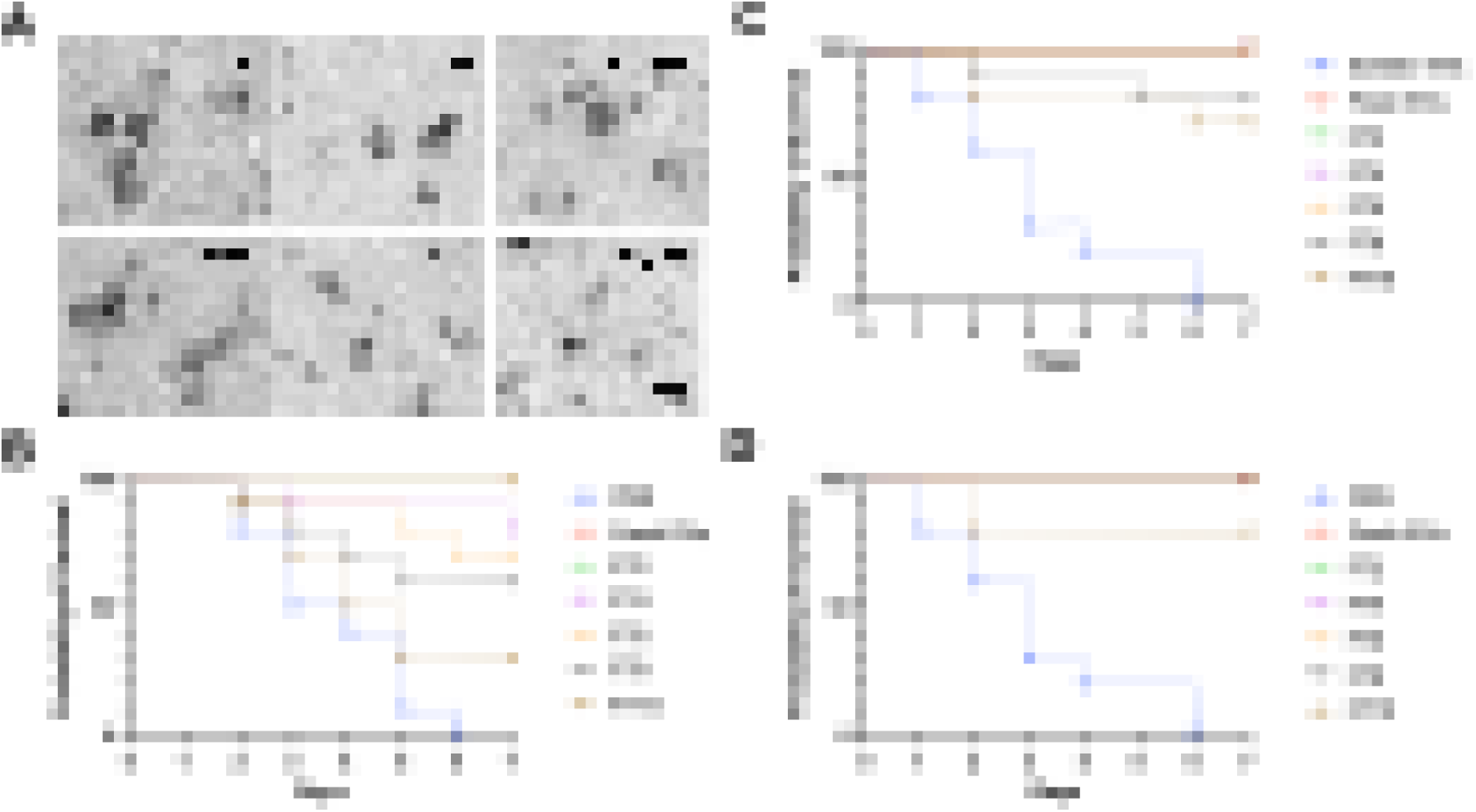
– Freeze-thaw cycles reduce the protective effect of *C. albicans* EVs. EVs released by *C. albicans* ATCC90028 were stored at -80 °C for a week and then consecutive cycles of freeze-thaw (FT) were performed on aliquots for 2, 4, 6, 8 or 10 cycles. (A) EVs were negatively stained with contrast and then visualized by TEM (CT, control at -80 °C). (B, C and D) Survival curves of *G. mellonella* larvae pretreated with fresh EVs submitted to 2, 4, 6, 8 or 10 FT cycles in the absence (B) or presence (C and D) of 10% sorbitol followed two days later by a lethal inoculum of *C. albicans* yeasts. In D, the sorbitol was removed after ultracentrifugation. Survival was monitored for 7 days (n=10). The survival differences between each EV groups and PBS were analyzed by individual paired Log-rank (Mantel-Cox). Results are representative of two independent experiments. Bars, 200 nm. * p= 0.029 and ** p= 0.0013.

### Biological activities of VCRT-EVs were similar to *in natura* EVs storage at -80 ⁰C

Although no statistical differences in recovery were observed among the different storage conditions for the EVs, the trend toward higher recovery in the VCRT and VCLT groups led us to select VCRT for evaluating their ability to protect insects against *C. albicans* infection. As anticipated from the physical analysis, suspended VCRT-EVs retained their capacity to protect *G. mellonella* larvae (Fig. 8). Both *in natura* stored EVs (EVs RT, EVs -4 °C and EVs -20 °C) and dried VCRT-EVs, (VCRT RT, VCRT -4 °C, VCRT -20 °C), demonstrated similar effectiveness in insect protection. Even when stored at room temperature, there was only a slight reduction in protection, which was not statistically significant, confirming the stability of these compartments even after storage under the diverse conditions employed in our experiments.

### Freeze-thaw cycles impacted the immunoactiviy of EVs from *C. albicans*

After every two cycles of freeze-thaw, *C. albicans* EVs were suspended in PBS, negatively stained with contrast and then visualized by TEM. EVs similar to control conditions were observed independent of the number of cycles, but the ability of *C. albicans* EVs to protect larvae from *G. mellonella* was significantly impacted by the number of freeze-thaw cycles (Fig 9B). Our experiments demonstrated that after 4, 6, 8 or 10 freeze-thaw cycles the protective effect was reduced by approximately 20, 30, 40 and 70%, respectively. Previous addition of sorbitol effectively preserved the protective effect of the EVs, which was otherwise compromised by freeze-thaw cycles. Importantly, this protective effect was observed both in the presence of sorbitol and following its removal via ultracentrifugation, yielding comparable results (Fig 9C and D).

## Discussion

In a previous study using an Ultrafiltration system as part of a protocol to isolate fungal EVs, we demonstrated that *C. albicans* EVs stored for a week at temperatures of 4, -20, or -80 °C exhibited diameters comparable to fresh preparations (4). Moreover, these EVs retained their antigenic properties and effectively protected insects against lethal challenges with *C. albicans* yeasts, affirming the potential utility of fungal EVs for vaccine formulations. However, extended storage can compromise certain biological preparations such as vaccines, possibly impacting their efficacy for immunization. According to our data, the size of *C. albicans* EVs showed major fluctuations after 4 and 5 years of storage as we detected an increase in the number EVs with higher diameters, which may be explained by EVs aggregation or fusion, a phenomena recently reported during storage of EVs from murine microglia (21). Although statistical differences were observed between the sizes of fresh EVs and EVs stored for up to 3 years, these EVs were similar in their size distributions, indicating that the EVs stored for 3 or fewer years underwent only small changes in size.

In the context of vaccines, the potency serves as a stability-indicating parameter, reflecting how environmental conditions can influence the vaccine’s immunogenicity and its subsequent protective effectiveness. We undertook a comparison between the effectiveness of long-term stored EVs and freshly prepared EVs using the *G. mellonella* candidiasis model. Surprisingly, all EVs protected the insects. However, the fresh EVs displayed the higher efficacy. Together, these results indicated that EVs from *C. albicans* may aggregate or fuse after long-term periods of storage, but they do not lose their biological capacity under the different conditions examined.

Advances in EV research field have enabled the development of different isolation protocols, improving the recovery efficiency and quality of samples. Initially, the ultrafiltration systems were employed as the primary method for isolating fungal EVs, and most of our previous investigations utilized this approach (4, 6, 14, 22). However, this protocol was time-consuming for large supernatant volumes and certain fungal species posed an additional challenge due to the high viscosities of the culture medium following fungal growth, leading to a significant reduction in the recovery of EVs. This challenge was particularly observed with *C. neoformans* where isolation of EVs is markedly compromised by the production of the yeast cell polysaccharide capsule.

To address these issues, Reis and colleagues (20) have developed a simplified isolation protocol using solid medium for cultivation of fungal cells. This method reduces the time required for the process by starting with fungal cultures and ending with ultracentrifuged EVs. This protocol also appears to be the most cost-effective method as it does not require filtration systems for supernatant concentration. An additional advantage of this protocol is its versatility in isolating EVs from various species and strains, making it suitable for screening studies. More recently, the utilization of tangential flow ultrafiltration has facilitated even faster isolations of EVs (19). This technique enables the filtration of larger volumes of medium within a few hours, resulting in the recovery of substantial quantities of EVs and has been used for isolating fungal EVs (23). The tangential flow system yielded superior results in a shorter timeframe. While the costs associated with purchasing both filtration systems for liquid medium are similar, it is worth noting that a washing step should be factored in for the tangential flow system when cassettes are reused.

In the present study, we found that EVs isolated from solid medium cultures displayed a more diverse size profile, with a broader range in diameter, compared to cells isolated by flow-based systems. Since the EVs in solid medium have a small area to diffuse it is possible that their constant contact promoted their fusion or agglutination. However, we cannot rule out that EVs with distinct sizes are produced in this culture condition. Additional studies are necessary to determine whether the global composition of these EVs are distinct. Previous studies from our lab demonstrated that EVs released by *C. albicans* yeast cells grown on solid medium displayed the same yeast-to-hyphae inhibitory activity as EVs obtained from liquid culture medium (14). Furthermore, independent of the isolation method used, the *C. albicans* EVs kept their biological properties, protecting the insects against a lethal challenge with *C. albicans*. Given that the tangential flow system generated the best EVs yields for larger volumes of medium we continued our experiments using this protocol for EVs isolation.

Various metrics, including the number of EVs, RNA levels, protein concentrations, lipid content and even the quantification of specific molecules have been used to assess various EVs characteristics. Indeed, any of these analysis should be sufficient for experiment normalization and comparison when the level of EVs purity is high. However, obtaining large yields of samples with high lelvel of purity remains one of the major difficulties in the EVs field. In this context, applicability of EVs faces challenges due to potential batch-to-batch variations can be a potential confounder to EV research. In situations where low-specificicity methods are employed to isolate EVs, multiple measurements are required for assurance of experimental validity (1). Previous work from our group indicated that the couting methods currently used to study *C. albicans* EVs are not precise, since the amount of small EVs are usually underestimated (24). In our prior experiments we observed that distinct *Candida* strains carry different amounts of lipids and proteins, reinforcing findings demostranting that different strains may produce EVs with distinct cargo (9). However, we also now demonstrated that the sterol-protein ratio normalized by the number of cells could be used as a valid strategy to incorporate as a quality control step for fungal EVs isolation.

Freeze-drying is a widely used method for preserving antigen properties in vaccine formulations (25). However, when the water content decreases, lipid membranes transition from a liquid crystalline to a gel-like state, potentially causing phase separation and membrane-bound EVs disruption and leakage during freezing drying or rehydration (26). Earlier studies shown that cryoprotectants like trehalose, sucrose, glucose, fructose and sorbitol can maintain the physical attributes of nanoparticles (21, 27, 28). However, unlike human EVs, fungal EVs possess glycan and polysaccharide coatings (11), which may add an extra layer of protection during freeze-drying processes. In our experimental conditions, *C. albicans* EVs that were dried using vacuum centrifugation at room temperature or -4 °C, stored for a week at 4 °C or -20 °C, and then rehydrated, maintained their physical stability, displaying similar structures as fresh EVs. Notably, differences emerged when the dried EVs were kept at room temperature. Under these circumstances, EVs exhibited a sligh increse in their sizes. Our findings emphasized the significance of temperature during *C. albicans* EV storage. However, while some EVs might have been disrupted in the experiments, *G. mellonella* treated with these EVs were still robustly protected against *C. albicans* infection. Collectively, these results imply that the protective effects were not solely reliant on complete EV integrity. Further investigations in murine models are warranted to explore potential toxic effects of these altered EVs. We also evaluated the stability of the EVs after exposure to the different conditions evaluated here, to determine the yields of recovery of these compartments under each experimental setup. A substantial loss of EVs was observed after a single ultracentrifugation step, with proportional decreases in protein and lipid content. Notably, complete EV disruption after 24 h in detergent suggested the absence of significant protein-based precipitates, a common contaminant specially in EV preparations using ultracentrifugation protocols (29, 30). Although no statistically differences in recovery were detected across the distinct drying and storage conditions, EVs dried at lower temperatures tended to yield higher recovery rates. Future studies employing high-sensitivity instrumentation will be important to precisely quantify the recovery in particle numbers.

In our earlier studies, EVs were either used immediately as fresh preparations or prepared in aliquots from a single thaw, with any remaining EVs discarded. When we examined the impact of freeze-thaw cycles, a notable reduction in protective activity was observed after four cycles. Since no additional ultracentrifdugation step was added after the freeze-thaw cycles we hypothesized that the antigenicity was lost most probably due to structural changes in the proteins carried by the EVs. Despite our TEM analysis indicating the presence of intact EVs, suggesting physical preservation, we cannot dismiss the possibility of membrane disruption and fusion. Our results suggest that sorbitol can effectively serve as a cryoprotectant for EVs undergoing a limited number of freeze-thaw cycles, but further investigations are imperative to elucidate the mechanisms by which EVs lose their antigenicity under repeat freezeing and thawing. As no protective effect was observed following treatment of larvae with sorbitol, we conclude that, under the conditions used in our experiments, this sugar alone was not sufficient to elicit an innate immune response.

EVs released by pathogens have emerged as promising vaccine formulations due to their ability to compartimentalize and concentrate a diversity of native immunogens, including Pathogen-Associated Molecular Patterns (PAMPS) and other bioactive compounds (5, 31). In the context of vaccine development, ensuring the stability of EVs formulations is crucial. Here, we demonstrated that it is possible to mantain physical and antigenic properties of *C. albicans* EVs under long-term storage and vacuum-drying. Remarkably, while variations in the sizes of EVs were noted according to their storage conditions, their established functions were largely unchanged, with the exception of EVs subjected to repeated freezes and thaws. Together, our data support the idea that fungal EVs should be further explored and exploited as vaccine formulations to combat fungal infections.

## Acknowledgments.

This work was supported by grants from the Brazilian agency Conselho Nacional de Desenvolvimento Científico e Tecnológico (CNPq, grants 408711/2017-7 to L.N.), FAPERJ (E-26/202.809/2018 and E-26/201.032/2022 to LN), FIOCRUZ (grants VPPCB-007-FIO-18 and VPPIS-001-FIO18) and Coordenação de Aperfeiçoamento de Pessoal de Nível Superior (CAPES, Finance Code 001). J.J.A.B was supported by FAPERJ (Fundação de Amparo à Pesquisa do Estado do Rio de Janeiro, Processo SEI E-26/204.281/2021).. LN is supported by Rede Micologia FAPERJ (E-26/211.300/2021). MLR was funded by the National Institute of Allergy and Infectious Diseases of the National Institutes of Health under Award Number R01AI183314, as well as CNPq grants 402651/2024-3, 404365/2023-0 and 304998/2022-2, the Program for Research Stimulation (PEP) of the Carlos Chagas Institute of Fiocruz, and the FAILSAFE/Global AMR Innovation Fund (GAMRIF) initiative, grant FR1-49 The authors declare no conflict of interest.

